# Morse-clustering of a Topological Data Analysis Network Identifies Phenotypes of Asthma Based on Blood Gene Expression Profiles

**DOI:** 10.1101/516328

**Authors:** James P R Schofield, Fabio Strazzeri, Jeannette Bigler, Michael Boedigheimer, Ian M Adcock, Kian Fan Chung, Aruna Bansal, Richard Knowles, Sven-Erik Dahlen, Craig E. Wheelock, Kai Sun, Ioannis Pandis, John Riley, Charles Auffray, Bertrand De Meulder, Diane Lefaudeux, Devi Ramanan, Ana R Sousa, Peter J Sterk, Rob. M Ewing, Ben D Macarthur, Ratko Djukanovic, Ruben Sanchez-Garcia, Paul J Skipp

## Abstract

Stratified medicine requires discretisation of disease populations for targeted treatments. We have developed and applied a discrete Morse theory clustering algorithm to a Topological Data Analysis (TDA) network model of 498 gene expression profiles of peripheral blood from asthma and healthy participants. The Morse clustering algorithm defined nine clusters, BC1-9, representing molecular phenotypes with discrete phenotypes including Type-1, 2 & 17 cytokine inflammatory pathways. The TDA network model and clusters were also characterised by activity of glucocorticoid receptor signalling associated with different expression profiles of glucocorticoid receptor (GR), according to microarray probesets targeted to the start or end of the GR mRNA’s 3’ UTR; suggesting differential GR mRNA processing as a possible driver of asthma phenotypes including steroid insensitivity.

## Introduction

Asthma is ranked 16^th^ among the leading causes of years lived with disability and affects 339 million people worldwide. Asthma is characterised by an expiratory airflow limitation, typically reported as forced expiratory volume in one second (FEV_1_). Treated is primarily with β2-agonists which relax airway smooth muscle, and corticosteroids which reduce underlying inflammation. Drugs have also been developed to target specific inflammatory pathways such as the T2 biologics, which reduce asthma exacerbation frequency by around 50%^1,2^. Improved understanding of asthma disease progression and molecular sub-phenotypes should improve the use and development of new targeted therapeutics. In this study, we used data from the U-BIOPRED (Unbiased BIOmarkers for the Prediction of respiratory disease outcomes) project, the largest multi-centre asthma programme to date, involving 20 academic institutions, 11 pharmaceutical companies and patient groups and charities, with the aim to improve understanding of the complex molecular mechanisms underpinning asthma and identify useful biomarkers^3–10^.

Asthma is characterized by variability in symptoms and treatment response. Around half of asthma is thought to arise from T-2 immunity, driven by IL4, IL5 and IL13 cytokine associated with recruitment of eosinophils into airways^11^. Additionally, high sputum neutrophil counts are associated with reduced post-bronchodilator FEV_1_^12^. Corticosteroids are routinely used to reduce airway inflammation in asthma by activating glucocorticoid receptor (GR) and suppressing NF-κB activity which regulates expression of pro-inflammatory cytokines and cyclo-oxygenase 2 (COX2) as well as inducible nitric oxide synthase (iNOS). However, patients with severe asthma, particularly T-2-low and T-17-high asthma^13^, respond poorly to corticosteroids, but it is not known why. The relative expression of GR-α and GR-β protein isoforms, resulting from alternative splicing, influences steroid insensitivity, as GR-β does not bind GC and inhibits GR-α activity by forming a heterodimer^14^. GR protein expression is further regulated by ARE-mediated degradation of GR mRNA targeting the AU-rich elements within the 3’ UTR^15^.

Topological Data Analysis (TDA) is an unsupervised machine learning tool suitable for analysis of high-dimensional datasets^16,17,18^. Application of TDA via the Mapper algorithm generates a TDA network model, a compressed representation of high-dimensional data with major features embedded where similar data points are grouped into nodes, and nodes with common data points are connected by edges. We have previously reported an analysis of differentially expressed genes (DEGs) from gene expression profiling of 498 gene expression profiles of peripheral blood from participants in the U-BIOPRED (Unbiased Biomarkers in Prediction of Respiratory Disease Outcomes) study^10^. Unbiased hierarchical clustering of DEGs identified two sub-groups, one enriched for patients with severe asthma, use of oral corticosteroids and blood neutrophilia, and a second cluster composed of mixed-severity asthmatics and healthy individuals. We generated a Topological Data Analysis (TDA) network model of the same gene expression data using the Ayasdi TDA software platform and found these two clusters represented by different regions of the TDA network model. In this study, we investigated the continuous variation of clinical and molecular biology in the TDA network model representing the shape of asthma disease pathology; shedding light on possible routes of disease progression.

Stratification of disease allows targeted treatment for improved patient outcome, so we developed and applied a Morse-clustering algorithm to discretise the continuous TDA network model of patients into clusters representing different molecular phenotypes of asthma sub-types. Clusters within TDA networks have typically been delineated by eye^18,19,20^, without algorithmic reproducibility and few studies have used the standard network clustering algorithm, community clustering, via the Ayasdi Python SDK. The community clustering algorithm is limited as it only analyses connectivity between nodes without considering the density of data points clustered within nodes, an important dimension in TDA network models. This 3^rd^ dimension in the TDA network can be visualised by colouring (Fig. 3A & B) and the TDA network can, therefore, be considered as a connected 3D map of data points clustered around peaks that represent conserved sub-types or phenotypes of major features, which in the study of patient gene expression reflect biological pathway modulations underlying disease phenotypes. Discrete Morse theory relates the flow (gradients) on a discrete object, such as a network, with its topology^21^. Here we apply Morse theory to measure the gradients and connected peaks within a TDA network, thus delineating clusters according to key features of the dataset. We have developed a Python script to apply Morse-based clustering of TDA networks in the open source Mapper TDA software or through the Ayasdi Python software development kit (SDK) which we believe will add value to future analyses. This Morse-clustering algorithm identified nine clusters, BC1-9, representing discrete molecular phenotypes characterised by differences in circulating immune cell populations, activation of T-1, -2 & -17 cytokine inflammatory pathways, and the activity of glucocorticoid receptor signalling and novel differences in glucocorticoid receptor mRNA isoforms.

**Figure 1.**
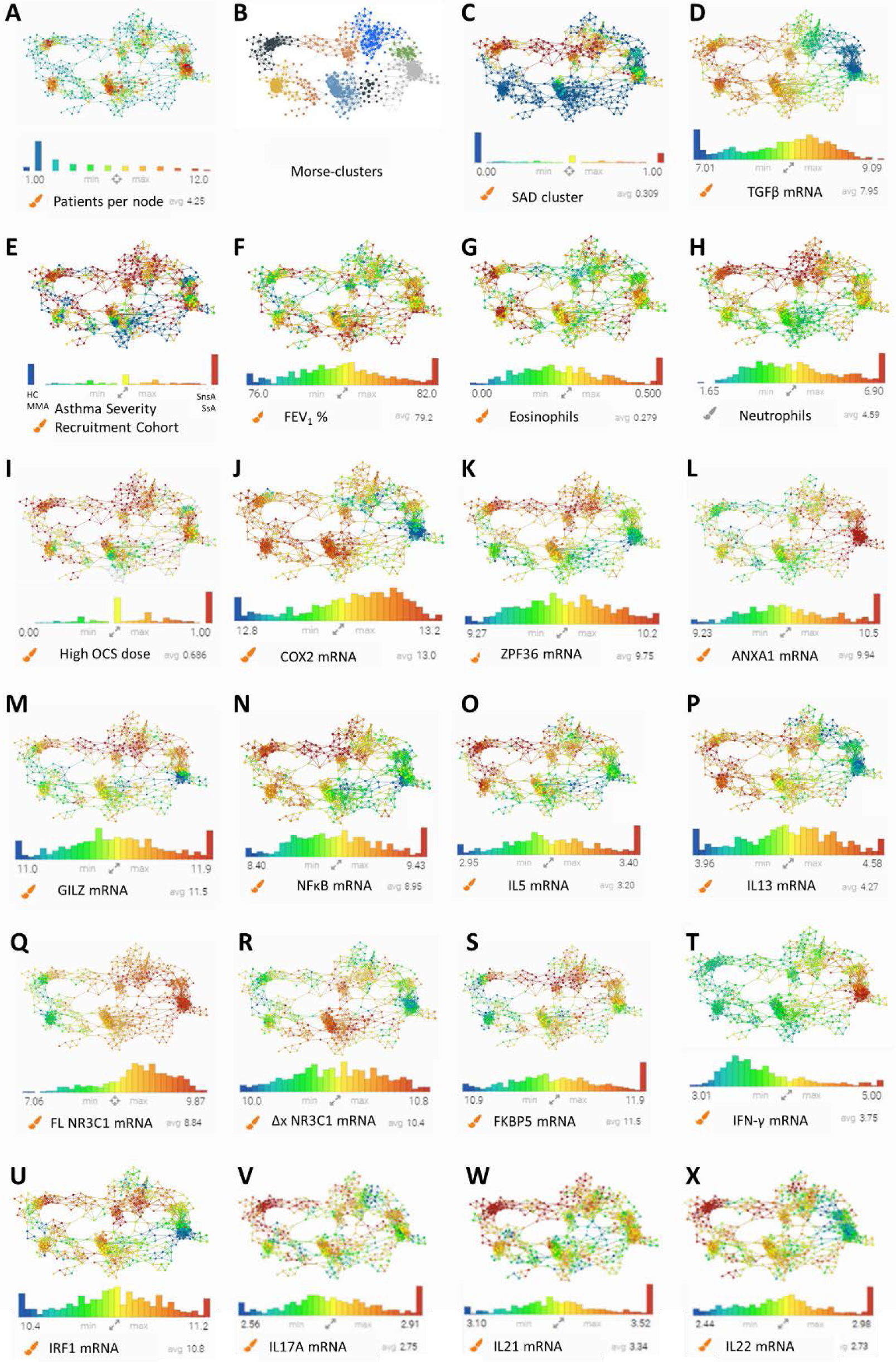
Selected gene expression distribution a cross the TDA network. Selected gene expression distribution across the TDA network. Colours in legends denote the concentrations of the gene expression, ranging from blue (low) to red (high).

**Figure 2.**
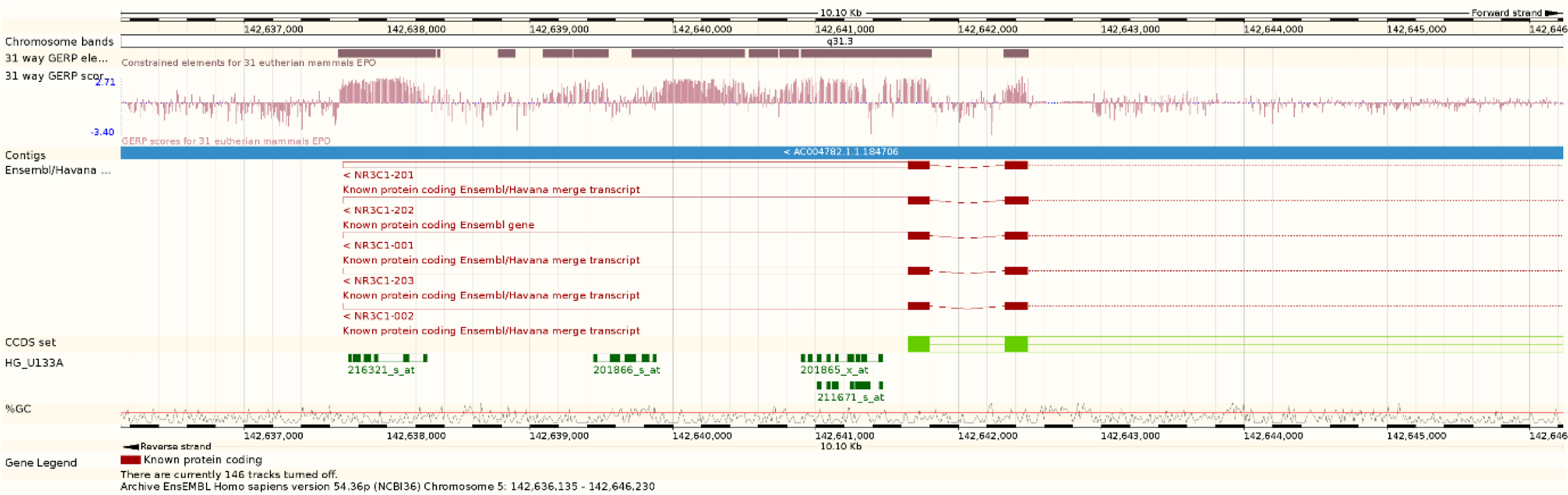
The chromosome binding locations of the Affymetrix NR3C1 probes. The binding locations of the Affymetrix NR3C1 probes and corresponding NCBI RefSeq sequences aligned to the Human genome. NR3C1 probesets 201865_x_at and 211671_s_at target isoforms with truncated 3’ UTR: Δx NR3C1. Probesets 201866_s_at and 216321_s_at target NR3C1 mRNAs towards the end of the 3’ UTR annotated in the RefSeq genes. Image generated using the Ensembl Genome Browser: https://genome.ucsc.edu

**Figure 3.**
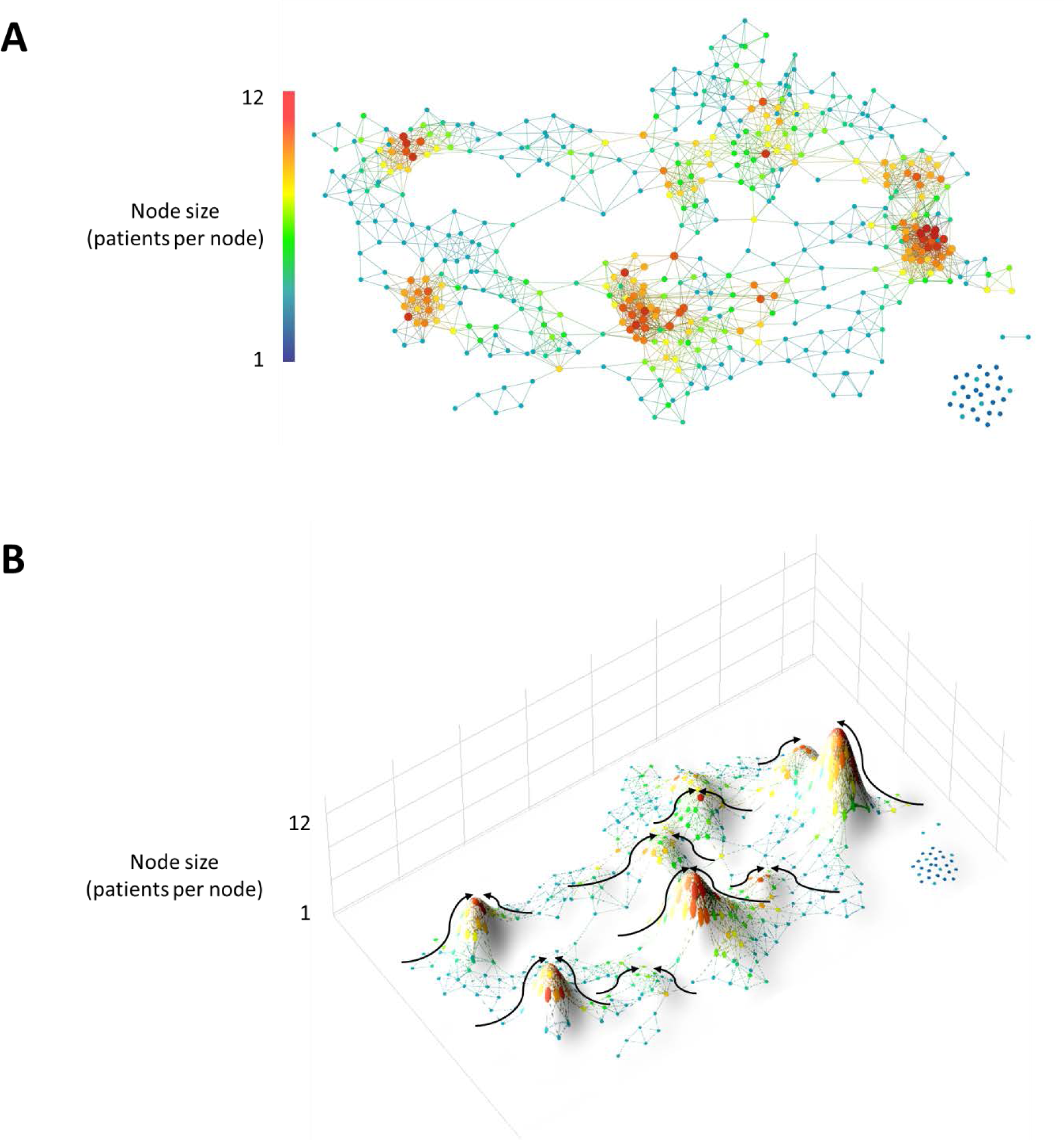
Morse-clustering of the TDA network of UBIOPRED gene expression profiling of peripheral blood. TDA network landscape of correlated gene expression (54,613 probesets, n = 498). Metric: norm correlation. Lenses: neighbourhood lens 1 (resolution, 100 bins; gain, ×6), neighbourhood lens 2 (resolution, 100 bins; gain, ×6) **(A)**. The vector (node value) is a 3^rd^ dimension in TDA networks, in a standard heatmap colouring of a TDA network, the colour represents the 3^rd^ dimension **(B)**. Arrows indicate the gradients of the 3-dimensional topology measured by Morse-based clustering identifying the ‘peaks’ as clusters of subjects with similar profiles of analysed variables.

## Results

The TDA network model of peripheral blood gene expression from 498 participants in the U-BIOPRED asthma study consisted of a hub with an increased prevalence of healthy participants and connected flares with increased prevalence of severe asthma and decreased FEV1, reflecting multiple interconnected possible routes of disease progression (Fig. 1). Regions of the TDA network with highest eosinophil counts (Fig. 1G) had high prevalence of severe asthma (Fig. 1E) and were associated with high COX2, NF-κB, IL5, IL13 (Fig. 1J, N, O, P), and low IFN-γ and GR mRNA (Fig. 1T, Q, R). There was a distinct pattern across the TDA network model of GR mRNA expression according to probesets targeting the start of the 3’ UTR (probesets 201865_x_at and 211671_s_at, illustrated as Δx NR3C1 mRNA in Fig. 1R) and a different pattern according to probesets targeting towards the end of the 3’ UTR (probesets 201866_s_at and 216321_s_at, illustrated as FL NR3C1 mRNA in Fig. 1Q). The binding locations of the Affymetrix NR3C1 probes and corresponding NCBI RefSeq sequences are shown mapped onto the Human genome in figure 2. We hypothesized that the Δx NR3C1 mRNA has a truncated 3’ UTR compared to the FL NR3C1; meaning Δx NR3C1 has fewer AU-rich elements (AREs), and is missing a miR 486 target sequence, compared to the FL NR3C1 mRNA. The TDA network was polarised by FL NR3C1 (Fig. 1Q) and associated GR-responsive genes, COX2, ANXA1 and IFNγ (Fig. 1J, L, T). Probesets targeting the start of the 3’ UTR of GR mRNA indicated a different pattern of expression across the TDA model (Fig. 1R) and corresponded to OCS dose (Fig. 1I) and GR-responsive gene expression, ZPF36, GILZ, FKBP5 (Fig. 1K, M, S).

To define groups of people with similar gene expression signatures from the TDA network model, we developed and applied a Morse-clustering algorithm. The Morse-clustering algorithm identified 9 clusters which we termed BC1 to 9. The reporter operating characteristic (ROC) area under the curve (AUC) for the 9 clusters ranged from 0.76 to 0.97, representing very good to excellent prediction of cluster classification in the test set based on a logistic regression model identifying predictors of the cluster in the training set (Fig. 4). BC1-9 were found to have activation of cytokine-mediated inflammatory pathways consistent with their distribution on the TDA network model with trends identified in pathway and upstream regulator activation across the clusters (Table1 & 2). BC1 was predominantly severe asthmatics, with reduced lung function, represented by low FEV_1_. BC1 also had a T-17 signature of gene expression^22^, with increased expression of IL17A, IL21 and IL22 (q = 1.31E^-5^, 7.99E^-4^, 1.71E^-3^). BC1 had decreased expression of β-2 adrenergic receptor (ADRB2) mRNA the protein product of which is involved in smooth muscle relaxation and bronchodilatation. Cystatin D (CST5) was predicted as the most activated upstream regulator of gene expression in BC1 but was also highly activated in BC9 and 8 (Table 2).

**Table 1.** Molecular pathways enriched in the 9 clusters. IPA identified significantly enriched (p<0.05) canonical pathways of gene expression in clusters (the top 5 pathways for clusters BC1-9 are shown). Values are z-scores, reflecting both the enrichment of specific transcription factor-regulated genes in the pathways and the degree of activation/inhibition. The z-scores are coloured blue (greatest downregulated transcription factor-regulated gene expression) to red (greatest upregulated transcription factor-regulated gene expression).

| Canonical Pathway | Sub-phenotype |  |  |  |  |  |  |  |  |
| --- | --- | --- | --- | --- | --- | --- | --- | --- | --- |
|  | BC1 | BC9 | BC8 | BC2 | BC7 | BC3 | BC6 | BC4 | BC5 |
| Serotonin Degradation | 3.1 |  |  |  |  |  |  |  |  |
| Superpathway of Melatonin Degradation | 2.5 |  |  |  |  |  |  |  |  |
| Melatonin Degradation I | 2.5 |  |  |  |  |  |  |  |  |
| Glutamate Receptor Signaling | 2.4 |  |  |  |  |  |  |  |  |
| Neuropathic Pain Signaling In Dorsal Horn Neurons | 2.4 |  | -0.9 | -1.8 | 0.0 |  | -0.6 |  |  |
| Oxidative Phosphorylation |  | 3.5 | 3.5 |  | 4.0 | -4.4 |  | -4.1 |  |
| Glycolysis I |  |  | 3.0 |  | 2.8 | -1.9 |  | -2.5 |  |
| Role of p14/p19ARF in Tumor Suppression |  |  | 1.4 | 0.0 | 0.3 | -0.3 |  | 0.5 | -0.9 |
| Cyclins and Cell Cycle Regulation |  | 2.1 |  |  |  |  |  |  |  |
| TNFR1 Signaling |  | 1.9 | 1.7 | 0.9 | 2.1 | 0.3 | -1.6 | -2.2 | -0.3 |
| tRNA Charging |  | 1.4 | 2.7 |  | 3.1 | -2.7 |  | -1.6 |  |
| Gluconeogenesis I |  |  |  |  |  | -1.1 |  | -1.7 |  |
| iNOS Signaling |  | 0.8 | 2.3 | 3.3 | 3.5 | 3.1 | -2.5 | -2.2 |  |
| Toll-like Receptor Signaling |  |  |  | 3.2 |  | 3.5 |  |  |  |
| Type I Diabetes Mellitus Signaling |  | 1.0 | 2.1 | 3.0 | 3.3 | 2.4 | -2.6 | -2.9 | -0.4 |
| TREM1 Signaling |  |  |  | 2.9 |  | 3.7 |  |  |  |
| Neuroinflammation Signaling Pathway |  |  |  | 2.7 |  | 2.3 | -2.2 |  |  |
| IL-1 Signaling |  | -1.0 | -0.2 | 2.5 | 1.5 | 2.8 |  | -0.8 | 1.1 |
| Inflammasome pathway |  |  |  | 2.4 |  | 2.6 |  |  |  |
| D-myo-inositol (1,4,5,6)-Tetrakisphosphate Biosynthesis |  | -3.0 | -0.3 |  | 0.0 | 0.9 | 2.8 | 0.3 | 4.4 |
| D-myo-inositol (3,4,5,6)-tetrakisphosphate Biosynthesis |  | -3.0 | -0.3 |  | 0.0 | 0.9 | 2.8 | 0.3 | 4.4 |
| 3-phosphoinositide Biosynthesis |  | -3.8 | -0.7 |  | 0.5 | 0.2 | 2.7 | -0.3 | 4.5 |
| 3-phosphoinositide Degradation |  | -3.0 | -0.1 |  | 0.6 | 0.7 | 2.4 | 0.1 | 4.0 |
| Superpathway of Inositol Phosphate Compounds |  | -3.5 | -0.5 |  | 1.2 | 0.0 | 2.1 | -0.9 | 4.2 |
| Cell Cycle: G1/S Checkpoint Regulation |  | -1.7 |  |  |  | 0.6 |  | 2.0 | 1.4 |
| Antioxidant Action of Vitamin C |  | 0.0 |  | -0.7 |  | -0.9 |  | 2.0 |  |
| HIPPO signaling |  | 0.7 | -0.5 |  | -1.5 |  | 1.2 |  | 0.0 |
| Cardiac β-adrenergic Signaling | | -1.1 | | -2.2 | -1.0 | | | | |
| ERK5 Signaling |  | -3.3 | -1.3 |  | 0.2 | 1.1 | 1.8 | 1.6 | 2.0 |
| D-myo-inositol-5-phosphate Metabolism |  | -2.5 | -0.2 |  | 0.8 | 0.7 | 2.1 | 0.0 | 4.3 |

**Table 2.** Activated upstream regulators enriched in the clusters. Upstream regulators of gene expression (p<0.05) in clusters predicted by IPA (the top 5 upstream regulators for clusters BC1-9 are shown). Values shown are z-scores, reflecting both the enrichment of specific transcription factor-regulated genes in the pathways and the degree of activation/inhibition. The z-scores are coloured blue of varying intensity (greatest downregulated transcription factor-regulated gene expression) to varying red (greatest upregulated transcription factor-regulated gene expression).

| Upstream regulator | Sub-phenotype |  |  |  |  |  |  |  |  |
| --- | --- | --- | --- | --- | --- | --- | --- | --- | --- |
|  | BC1 | BC9 | BC8 | BC2 | BC7 | BC3 | BC6 | BC4 | BC5 |
| CST5 | 3.45 | 2.56 | 2.01 | 3.24 | 1.69 | 2.02 | -2.6 | -1.5 | -3.4 |
| TP63 |  |  |  |  |  | 1.79 | 0.17 |  |  |
| HSF1 |  |  | 1.31 |  |  | 2.13 |  |  |  |
| TGM2 |  |  |  | 5.91 |  | 3.85 | -4.4 |  |  |
| ERG |  | -1.6 |  | -0.3 |  |  | -1.4 |  | -0.9 |
| TAL1 |  | 3.31 | 2.42 |  |  |  |  |  |  |
| miR-486-5p (and other miRNAs w/seed CCUGUAC) |  | 2.91 |  | 0.37 | 1.33 | -1.2 | -3.3 | -2 | -2.6 |
| mir-486 |  | 2.89 |  | 0.24 |  | -1.2 | -3.3 | -2.1 | -2.6 |
| NUPR1 | 0.76 | 2.86 |  | 2.98 |  | 2.54 |  |  |  |
| RAE1 | 1.34 | 2.83 |  | 0.45 |  |  | -1.9 |  |  |
| SPP1 |  | 2.37 |  |  |  | -2.2 |  |  |  |
| TFEB |  |  | 2.98 |  |  |  |  |  |  |
| IL15 |  | 1.15 | 2.67 | 1.22 |  | -0.8 | -1.3 | -1.5 |  |
| miR-30a-3p (and other miRNAs w/seed UUUCAGU) |  | 2.82 | 2.63 | 1.63 |  |  | -1.3 | -1.6 | -2.2 |
| EIF2AK2 |  |  |  | 3.05 |  | 1.44 |  |  |  |
| CEBPA |  |  |  | 2.77 |  | 2.8 |  |  |  |
| PCGEM1 |  |  |  |  | 2.28 | -1.2 |  | -1.4 |  |
| LINC01139 |  |  |  | 1 | 2.24 | 0.45 |  |  |  |
| PLA2R1 |  | 1.25 | 1.04 |  |  |  |  | -1.6 |  |
| LDL |  |  |  | 1.39 |  | 1.93 |  |  |  |
| PPRC1 |  |  |  |  |  | 3.46 |  |  |  |
| PDGF BB |  |  |  |  |  | 3.31 |  |  |  |
| TNF |  |  |  |  |  | 3.11 |  |  |  |
| IL5 |  |  |  | 1.26 |  |  |  |  |  |
| CD24 |  | -5.3 | -5.2 |  | -3.9 | 1 | 4.41 | 4.67 | 5.11 |
| MYC | -2.9 | -2.6 |  | -4.5 |  | -2 | 3.06 | 0.74 |  |
| HELLS |  | -1 |  |  |  |  | 2.45 |  | 2.24 |
| MAPK1 |  |  |  | -2 |  |  |  |  |  |
| SAFB |  |  |  | -2.1 |  | -1.9 | 2.35 |  |  |
| SLC29A1 |  | -1.2 |  |  |  | 1.63 |  | 2.65 | 1.41 |
| WT1 |  |  | -1.6 | -1.1 |  | 1.61 | -0.2 |  |  |
| FSH |  | -2.1 | -2.3 | -0.4 |  | 0.43 | 1.96 | 2.62 | 2.72 |
| TCR |  |  |  | -0.7 |  | -0.8 |  | -1.8 | 2.49 |
| THOC5 |  | -2.2 |  |  |  |  | 1.63 |  | 2.45 |

**Figure 4.**
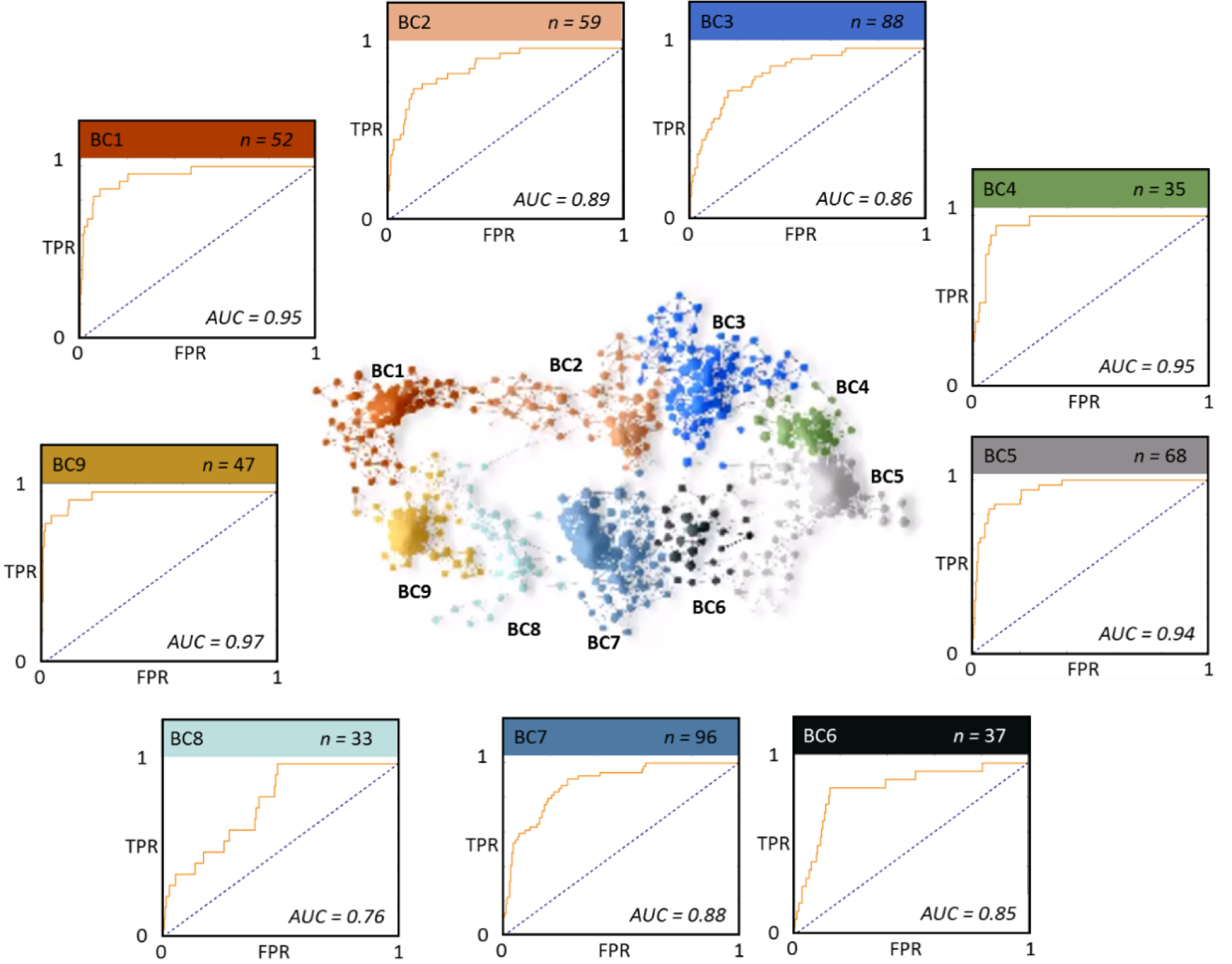
Clusters identified by Morse-clustering of the TDA network. **Centre:** TDA network coloured by clusters (BC1-9) identified using the Morse-based algorithm. **Outside:** Colour-coded ROC curves of cluster prediction success representative of cluster robustness.

## Discussion

The TDA network model identified familiar phenotypes of asthma and gave insight into potential routes of disease progression. For example, the furthest eosinophilic region from the ‘healthy hub’ was associated with high T-17 markers, TGFβ, IL17A, IL21, IL22 (Fig. 1D, V, W, X) and increased neutrophilia (Fig. 1H). The T-17 region was connected to the ‘healthy hub’ via the solely T-2 high region, suggesting disease progression from healthy to T-17 high via an only T-2-high phenotype. Differential expression of FL NR3C1 and Δx NR3C1 and corresponding expression patterns of GR-responsive genes suggests different functional responses to steroids across the TDA network model, associated with differential expression of GR mRNA isoforms.

The Morse-clustering algorithm identified 9 clusters, however, clusters BC4, 6 and 8 were small (n=35, 37, 33, respectively), with correspondingly low representation in the training and test sets which resulted in ROC curves whose shapes were not smooth and may have represented overfitting. The identified clusters represented groups of patients with significant differences in the activation of pathways related to inflammation, including pathways associated with glucocorticoid receptor (GR) signalling, Type (T)-2, T-1 and T-17 inflammatory responses. Transglutaminase (TGM2), a marker of T-2 inflammation^23^, was predicted in this study as the most activated upstream regulator of gene expression in BC2, 3, 7 and 8 (Table 2). It is known to catalyse the serotonin transamidation of glutamines (serotonylation), which regulates cell signalling and actin polymerization. BC2 and 3 were characterised by high TGM2-mediated gene expression, including Toll-like receptors (TLR) and iNOS signalling. TGM2 is also implicated in recruitment of eosinophils into asthmatic airways^11^, which was reflected in the highest sputum eosinophil count in BC2, but high sputum eosinophils counts were not seen in BC3 (Table 3). Melatonin, the end product of the serotonin pathway is a free radical scavenger, acting to suppress inflammation^24^. Pathways associated with tryptophan metabolism were enriched in cluster BC1; serotonin degradation was the most activated pathway identified by IPA (Table 1). Serotonin levels are known to be implicated in asthma pathology, and serum serotonin levels tend to be increased in patients with active asthma^25^. The increased activation of melatonin degradation in BC1 may contribute to the severe asthma phenotype.

**Table 3.** Clinical characteristics of the clusters. Clinical features associated with the TDA-defined asthma phenotypes. Values are shown as means and are colour coded on a heat scale for each variable; highest variable value is in red, lowest value in blue. FEV_1_: forced expiratory volume in one second (measured by spirometry). FVC: forced vital capacity. (%) Severe Asthma cluster (%) is the percentage of study participants previously identified in the severe asthma enriched cluster identified by hierarchical clustering^10^. ACQ5 or 7: asthma quality questionnaire consisting of 5 or 7 questions. AQLQ: asthma quality of life questionnaire. Sputum cells are shown as percentages of total inflammatory cells.

| Cluster | BC1 | BC2 | BC3 | BC4 | BC5 | BC6 | BC7 | BC8 | BC9 |
| --- | --- | --- | --- | --- | --- | --- | --- | --- | --- |
| Number of participants | 52 (10.44%) | 59 (11.84%) | 88 (17.67%) | 35 (7.02%) | 68 (13.65%) | 37 (7.42%) | 96 (19.27%) | 33 (6.62%) | 47 (9.43%) |
| FEV<sub>1</sub> (%) | 72.21 ± 24.64 | 66.04 ± 20.76 | 67.89 ± 25.16 | 76.06 ± 23.28 | 79.63 ± 23.62 | 87 ± 21.03 | 83.33 ± 23.83 | 78.57 ± 22.97 | 71.69 ± 24 |
| FVC (%) | 88.97 ± 20.9 | 88.83 ± 19.43 | 85.52 ± 23.42 | 95.12 ± 24.11 | 98.13 ± 21.01 | 99.77 ± 17.12 | 98.67 ± 19.96 | 95.12 ± 22.5 | 86.68 ± 21.35 |
| Severe Asthma (non-smoker) (%) | 69.2 | 50.8 | 38.6 | 42.8 | 33.8 | 43.2 | 18.7 | 51.5 | 51 |
| Severe Asthma (smoker) (%) | 9.6 | 23.7 | 21.5 | 17.1 | 19.1 | 10.8 | 15.6 | 18.1 | 17 |
| Mild-moderate Asthma (%) | 9.6 | 11.8 | 9 | 22.8 | 13.2 | 8.1 | 25 | 18.1 | 10.6 |
| Healthy (%) | 11.5 | 1.6 | 9 | 17.1 | 22 | 37.8 | 23.9 | 12.1 | 21.2 |
| Severe Asthma cluster (%) | 75 | 81 | 39 | 22 | 25 | 5 | 5 | 3 | 2 |
| Age | 51.44 ± 14.73 | 53.03 ± 14.44 | 51.07 ± 14.45 | 46.88 ± 16.45 | 44.07 ± 13.97 | 44.51 ± 14.87 | 45.22 ± 14.95 | 47.57 ± 15.47 | 50.8 ± 15.58 |
| Smoking (Pack Years) | 3.3 ± 11.44 | 6.38 ± 16.00 | 5.05 ± 11.69 | 3.64 ± 7.11 | 4.59 ± 10.87 | 2.66 ± 7.69 | 3.69 ± 10.56 | 5.07 ± 10.72 | 5.87 ± 14.52 |
| Mean ACQ5 | 1.69 ± 1.49 | 1.95 ± 1.23 | 1.83 ± 1.34 | 1.46 ± 1.51 | 1.44 ± 1.39 | 1.03 ± 1.41 | 1.18 ± 1.23 | 1.58 ± 1.36 | 1.65 ± 1.48 |
| Mean ACQ7 | 2 ± 1.65 | 2.31 ± 1.36 | 2.17 ± 1.5 | 1.66 ± 1.65 | 1.67 ± 1.52 | 1.15 ± 1.52 | 1.4 ± 1.37 | 1.82 ± 1.46 | 1.98 ± 1.61 |
| Mean AQLQ | 3.68 ± 2.24 | 4.64 ± 1.57 | 4.08 ± 2 | 3.6 ± 2.52 | 3.98 ± 2.35 | 3.16 ± 2.81 | 3.78 ± 2.59 | 3.74 ± 2.48 | 3.36 ± 2.24 |
| Admitted to ICU (%) | 0.25 ± 0.4 | 0.2 ± 0.54 | 0.17 ± 0.37 | 0.17 ± 0.17 | 0.23 ± 0.19 | 0.05 ± 0.13 | 0.13 ± 0.13 | 0.18 ± 0.18 | 0.17 ± 0.19 |
| Oral steroids (%) | 40.38 ± 46.57 | 54.24 ± 38.46 | 37.50 ± 40.45 | 17.14 ± 41.23 | 19.12 ± 39.79 | 13.51 ± 45.32 | 13.54 ± 44.21 | 18.18 ± 46.09 | 19.15 ± 44.31 |
| Blood periostin (ng/ml) | 46.57 ± 24.62 | 38.46 ± 23.24 | 40.45 ± 27.57 | 41.23 ± 27.09 | 39.79 ± 22.02 | 45.32 ± 24.13 | 44.21 ± 21.45 | 46.09 ± 19.88 | 44.31 ± 23.59 |
| Atopy (% positive) | 0.65 ± 29.81 | 0.67 ± 31.71 | 0.67 ± 32.66 | 0.68 ± 36.58 | 0.72 ± 31.78 | 0.56 ± 30.74 | 0.67 ± 33.75 | 0.66 ± 28.72 | 0.8 ± 26.34 |
| Exhaled NO (ppb) | 29.81 ± 22.04 | 31.71 ± 30.11 | 32.66 ± 26.52 | 36.58 ± 32.73 | 31.78 ± 30.61 | 30.74 ± 32.05 | 33.75 ± 31.02 | 28.72 ± 26.51 | 26.34 ± 14.71 |
| Blood eosinophils (x10<sup>3</sup>/μL) | 0.31 ± 0.3 | 0.18 ± 0.17 | 0.25 ± 0.28 | 0.21 ± 0.14 | 0.25 ± 0.25 | 0.23 ± 0.21 | 0.23 ± 0.2 | 0.29 ± 0.24 | 0.35 ± 0.33 |
| Blood neutrophils (x10<sup>3</sup>/μL) | 5.63 ± 2.3 | 6.78 ± 2.94 | 5.41 ± 2.35 | 4.35 ± 1.52 | 4.18 ± 1.86 | 3.32 ± 1.37 | 3.42 ± 1.09 | 3.99 ± 1.2 | 4.06 ± 1.75 |
| Blood lymphocytes (x10<sup>3</sup>/μL) | 2.06 ± 0.7 | 1.57 ± 0.7 | 1.83 ± 0.76 | 2 ± 0.47 | 1.91 ± 0.82 | 2.03 ± 0.73 | 1.87 ± 0.46 | 2.22 ± 0.66 | 2.14 ± 0.75 |
| Sputum Eosinophils (%) | 1.67 ± 5.16 | 6.37 ± 14.89 | 2.33 ± 9.42 | 1.77 ± 8.27 | 3.84 ± 12.49 | 5.79 ± 16.41 | 5.28 ± 12.42 | 4.47 ± 10.25 | 3.32 ± 12.41 |
| Sputum Neutrophils (%) | 30.18 ± 36.16 | 29.48 ± 34.25 | 5.7 ± 17.38 | 3.45 ± 12.12 | 17.37 ± 25.54 | 21.7 ± 28.31 | 28.65 ± 28.88 | 28.48 ± 29.83 | 24.83 ± 31.74 |
| Sputum Macrophages (%) | 13.65 ± 20.15 | 12.66 ± 17.96 | 3.16 ± 10.9 | 2.79 ± 9.48 | 17.89 ± 27.24 | 25.88 ± 33.44 | 30.62 ± 30.57 | 29.99 ± 30.84 | 26.38 ± 32.85 |
| Sputum Lymphocytes (%) | 0.62 ± 1.26 | 0.61 ± 1.06 | 0.15 ± 0.65 | 0.53 ± 2.29 | 0.57 ± 0.99 | 0.64 ± 0.84 | 1.04 ± 1.34 | 0.68 ± 0.95 | 0.74 ± 1.21 |

T-cell acute lymphocytic leukemia protein 1 (TAL1) was identified as the top upstream regulator of gene expression in BC9, together with miR-486, which has previously been identified as a potential marker of childhood asthma in plasma^26^ and a promoter of NF-κB activity^27^. Our analysis predicted CD24 as the most activated upstream regulator of gene expression in BC6, 4, and 5. CD24 can reflect activity of one of its key transcription factors, c-myc, whose expression is inhibited by CST5. BC5 had high expression of IFN-γ mRNA (Fig. 1T), indicative of a T-1 response; however, IFN-γ-mediated gene expression was not upregulated in this group (Table 3).

The shape of the TDA network and patterns of gene expression representative of differentially activated pathways reflected both corticosteroids use and expression of GR mRNA. Clusters BC1-3, mostly representing those of the Severe Asthma enriched cluster previously reported^10^ (Fig. 1C), had the highest percentages of patients on OCS (Table 3). These clusters were also characterised by enrichment for patients on high doses of OCS, but other clusters were also enriched for patients with high OCS dose; particularly cluster BC5 (Fig. 1I). We observed common patterns of gene expression under the control of glucocorticoid response elements (GRE) that were differentially expressed between clusters, although the patterns were not necessarily consistent between GRE genes. This suggests different types of steroid response between the clusters. We did not find GR-signalling as a top upstream regulator of gene expression using IPA, because there are two signatures of GR-signalling which are alternately up and down regulated in the TDA structure. The expression of GRE genes, glucocorticoid-induced leucine zipper (GILZ), FK506-binding protein 5 (FKBP5) and Tristetraprolin (ZFP36) (Fig. 1M, S and K) were similarly distributed across Morse-clusters high in neutrophilic clusters of the top of the TDA network, BC1, 2, 3 & 4 and higher in the predominantly healthy cluster, BC7. However, the expression of Annexin A1, a classical indicator of steroid response, was very differently distributed between clusters (Fig. 1L) and was significantly higher in BC5 when compared to the other patients (q = 2.3E^-10^). Serotonin degradation, which is interdependent on GR signalling, was identified as the top canonical pathway enriched in BC1 (Table 1). In clusters BC1-3, there was increased expression of the RNA-binding protein, tristetraprolin (TTP), a negative regulator of mRNA half-life, binding to AREs in the 3’ UTR of target genes (Fig. 1K). Since the expression of TTP is regulated by a GRE site, GR-signalling causes increased ARE-mediated mRNA decay.

BC1 had low expression of short (Δx NR3C1) and long (FL NR3C1) GR mRNA and low expression of steroid-inducible anti-inflammatory mRNAs ANXA1 (Fig. 1L), SOCS1 and high expression of pro-inflammatory COX genes (Fig. 1J). We detected mixed levels of GILZ and FKBP5 (Fig. 1M & S). There was moderate expression of DUSP1 mRNA, another marker of GR activity. In the clusters on the left side of the TDA network there was high expression of NUPR1 which increases expression of p38MAPK, a key regulator of asthma pathogenesis^28^. Additionally, NUPR1 is known to activate phosphatidylinositol 3-kinases (PI3K)^29^ which activate phosphoinositide pathways; inositol-related metabolism was highly upregulated in BC5 and 6, where the expression of phosphoinositol (PI) phosphatases was increased relative to health. Conversely, the expression of PI phosphatases was decreased when compared to health in BC8 and 9. Clusters BC5 and 6 showed increased expression of the enzyme which catalyses the dephosphorylation of 1D-myo-inositol (3)-monophosphate to myo-inositol, inositol-1 (or 4)-monophosphatase, when compared to health, whereas BC1, 7, 8 and 9 had decreased expression relative to health. It has previously been reported that <u>myo-inositol</u> is increased in animal asthma models following steroid treatment^30^, suggesting differential steroid responses between these clusters. In contrast to BC1, BC5 and 6 had gene expression profiles characteristic of low GR responses, as indicated by activation of CD24-mediated gene expression and inactivation of CST5-mediated gene expression. CST5 is activated by vitamin D receptor (VDR) expression^31^, whose expression is regulated by steroid-induced GR signalling^32^ (Fig. 5). The enriched expression of inositol pathways in BC5 and 6 provided further support of a low GR response. Contraction of airway smooth muscle is initiated by increased cytosolic calcium ions (Ca^2+^), so this may, in part, explain the reduced FEV_1_ seen in these clusters.

**Figure 5.**
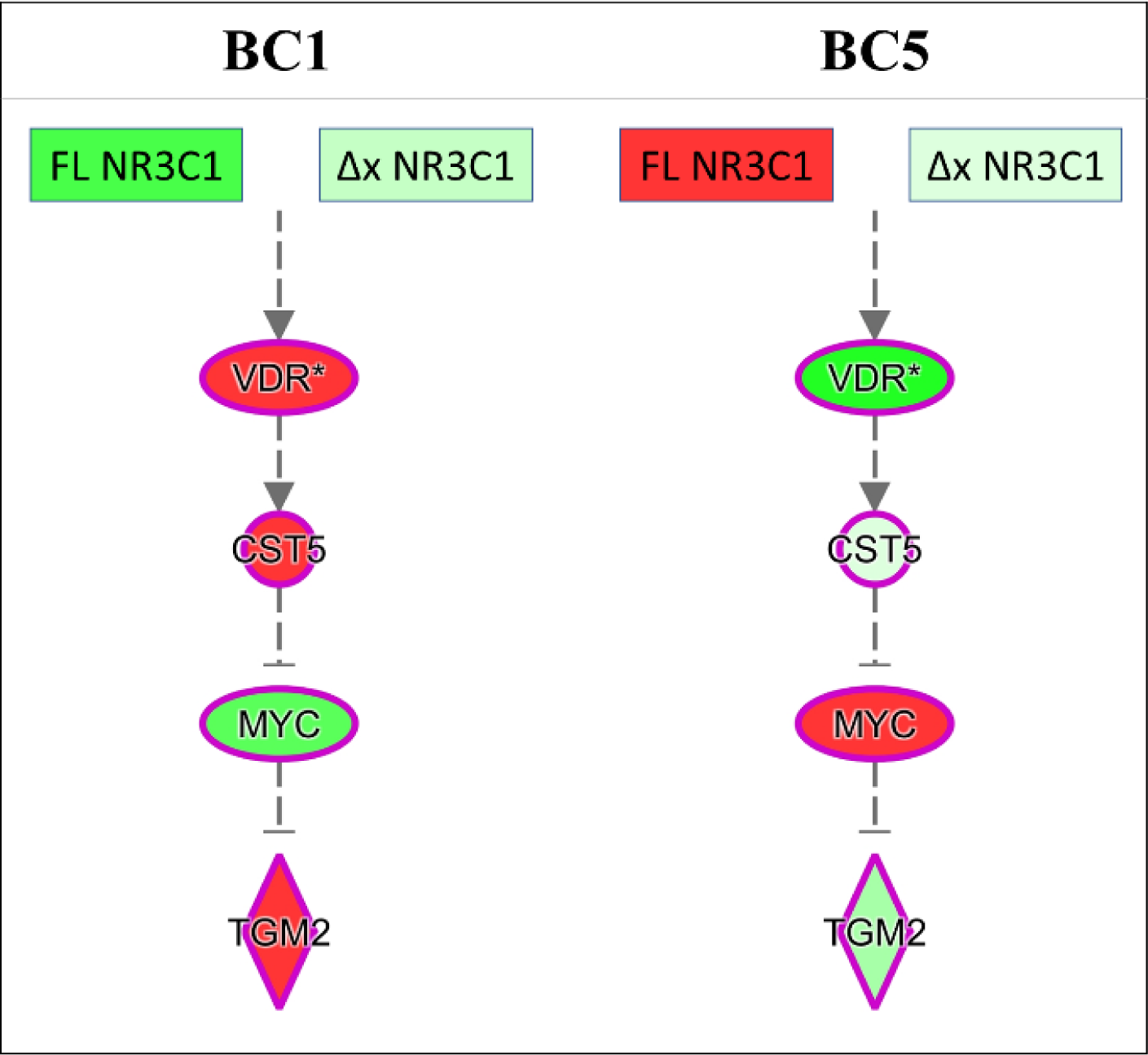
The regulatory gene pathway of NR3C1 transcript variants, and VDR, CST5, MYC & TGM2; identified as top upstream regulators by IPA (Table 2). Colours indicate gene expression relative to healthy participants, where green represents lower gene expression and red represents higher gene expression, white indicates no change (negative, positive and zero-fold change). Left column shows gene expression in cluster BC1, right column shows gene expression in BC5. Image generated using IPA.

We propose that Morse clustering can be applied to TDA networks of patient ‘omics data to identify sub-phenotypes of disease, thereby offering new insights into disease mechanisms and stratification of patients for more targeted drug development based on molecular mechanisms.

## Materials and Methods

### Study population

U-BIOPRED is a multi-centre prospective cohort study, involving 16 clinical centres in 11 European countries. Blood samples were analysed from 498 study participants; 246 non-smoking severe asthmatics, 88 smoking severe asthmatics, 77 non-smoking mild/moderate asthmatics and 87 non-smoking non-asthmatic individuals. It is registered on ClinicalTrials.gov (identifier: NCT01982162).

### Ethics Statement

The study was conducted in accordance with the principles expressed in the Declaration of Helsinki. It was approved by the Institutional Review Boards of all the participating institutions; Academic Medical Centre (AMC), Amsterdam; University Hospital Southampton NHS Trust; South Manchester Healthcare Trust; Protisvalor Méditerranée SAS; Karolinska University Hospital; Nottingham University Hospital; NIHR-Wellcome Trust Clinical Research Facility; and adhered to the standards set by the International Conference on Harmonization and Good Clinical Practice. All participants provided written informed consent.

### Microarray Analysis

RNA was isolated using the PAXgene Blood RNA kit (Qiagen, Valencia, CA) with on-column DNase treatment (Qiagen). RNA integrity was assessed using a Bioanalyzer 2100 (Agilent Technologies, Santa Clara, CA). Samples with RIN≥6 were processed for microarray as described (19) and hybridized onto Affymetrix HT HG-U133PM+ arrays (Affymetrix, Santa Clara, CA) using a GeneTitanR according to Affymetrix technical protocols. The microarray data are deposited in GEO under GSE69683.

### Training and Test Data Analysis Sets

The 498 samples available for analysis were randomized into training (n = 328) and validation sets (n = 170).

### Topological Data Analysis

#### Generating TDA graphs in Ayasdi Platform

The transcriptomics data were clustered by topological data analysis (TDA) as previously reported^10^, using Ayasdi Platform with a norm correlation metric and two Neighbourhood lenses. Correlation was measured using normalised values for the expression of each probeset (Metric: norm correlation). The space for clustering was generated using 100 bins in each dimension according to t-SNE -calculated vectors and 60% overlap between neighbouring bins (Fig 3A): two neighbourhood lenses, resolution = 100; gain, ×6).

#### Clustering of high patient density regions of TDA graphs

Using the Ayasdi TDA Platform, the magnitude of nodes was represented by a colour heatmap where the colour spectrum from blue to red represent the range from the lowest to highest levels. Discrete Morse theory was applied to cluster TDA nodes according to patient density. Data from each node’s neighbours were also used in calculating the annotation function, giving context to where a node lies within the broader topology, effectively ‘smoothing’ the data, decreasing noise and allowing identification of the most prominent peaks. To each node we assigned the annotation *f*: *V* → ℛ^2^ where for each node *CC*_*ii*_ we have

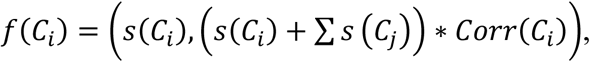

and *Corr*(*C*_*i*_) is the average correlation among all the patient in cluster-node *C*_*i*_. Differently from other clustering algorithms, as k-nearest neighbours, we do not assume that cluster-nodes with similar value with respect to *f* are similar, neither we expect that *f* is a kernel-based function which fits the data. Our approach instead assumes that *f* gives the cluster-nodes a hierarchical structure and the nodes’ connectivity is supplied by the Mapper network. In this way, with Morse, each cluster of nodes in the network has a structure of rooted tree and each leaf connects a cluster-node to a higher one (with respect to *f*) with the root the highest cluster-node.

#### Robustness of TDA network clusters evaluated by ROC analysis

We applied logistic regression to test the tightness of the clusters according to key features identified by logistic regression. A logistic regression model was trained on a pre-defined training set of (n = 328) and the classification accuracy tested on a test data set (n = 170). Accuracy of the logistic regression reflects reproducibility in the clustering, ie. robust classification assigned by clustering results in accurate classification of test data by an independently trained logistic regression model.

Affymetrix probes for NR3C1 were aligned with NCBI RefSeq genes using the Ensembl Genome browser 94.

#### Pathway analysis identified trends and discrete molecular features of clusters

The shape of data represented by a TDA network is defined by the lenses (t-SNE in this study), which are implicitly used as coordinates for plotting the network. These coordinates focus on differentially activated pathways because genes of a common pathway are more likely to be co-expressed, and patients are clustered by similarity in key features in a TDA network. Ingenuity pathway analysis (IPA) was used to identify pathways with enriched gene expression within each of the clusters (Table 1), many of which were activated in clusters neighbouring each other in the TDA network, reflecting a trend in the activation of key pathways across the TDA network.

## The U-BIOPRED Study Group

A. Bautmans^16^, A. Chaiboonchoe^13^, A. Mazein^13^, A. Sogbesan^17^, A. Meiser^4^, A. Menzies-Gow^17^, A. Berglind^18^, A.-S. Lantz^7^, A.J. James^8^, A. Petrén^8^, A.F. Behndig^19^, A. Dijkhuis^13^, A. Postle^20^, A. Rowe^21^, A. Vink^22^, A. Pacino^23^, A. Aliprantis^24^, A. Wagener^13^, A. Braun^25^, A. D’Amico^26^, A. Woodcock^27^, B. Smids^13^, B. Lambrecht^28^, B. Nicholas^20^, B. Nordlund^18^, B. Thornton^29^, A. Roberts^30^, B. Flood^30^, C. Mathon^31^, C. Smith^32^, C. Holweg^33^, C. Compton^10^, C. von Garnier^34^, C. Rossios^4^, C. Barber^15^, C.S. Murray^27^, C. Wiegman^4^, C. Schoelch^35^, C. Faulenbach^36^, C. Coleman^30^, C. Gomez^8^, D. Erzen^35^, D. Balgoma^8^, D. Gibeon^4^, D. Myles^10^, D. Supple^30^, D. Campagna^37^, D. Burg^1^, D.E. Shaw^38^, D. Staykova^20^, E. Bel^13^, E. Henriksson^39^, E. Yeyasingham^40^, E. Ray^32^, E.J. Kennington^30^, F. Singer^41^, F. Wald^35^, F. Baribaud^42^, G. Galffy^43^, G. Pennazza^26^, G. Santini^44^, G. Roberts^45^, G. Bochenek^46^, G. Hedlin^18^, H. Bisgaard^47^, H. Ahmed^11^, H. Gallart^8^, H. Knobel^22^, I. Horvath^43^, I. De Lepeleire^16^, I. Delin^8^, J. Musial^46^, J. Martin^32^, J. Versnel^30^, J. Hohlfeld^25^, J. Edwards^30^, J. Smith^30^, J.P. Carvalho da Purificação Rocha^17^, J. Kolmert^8^, J.G. Matthews^33^, J. Haughney^48^, J.-O. Thörngren^49^, J. Konradsen^18^, J. Thorsen^47^, J. Ward^50^, J. Brandsma^20^, J. Beleta^51^, J. De Alba^51^, J. Östling^52^, J. Vestbo^27^, J. Gent^53^, J. Corfield^54^, J. Kamphuis^55^, K. Tariq^56^, K. Strandberg^39^, A. Knox^57^, K.M. Smith^57^, K. Riemann^35^, K. Nething^35^, K. van Drunen^13^, K. Dyson^58^, K. Gove^15^, K. Russell^4^, K. Alving^59^, K. Bøonnelykke^47^, K. Fichtner^35^, K. Zwinderman^13^, K. Wetzel^35^, L. Ravanetti^13^, L. Larsson^60^, L. Pahus^61^, L. Metcalf^30^, L. Carayannopoulos^29^, L. Tamasi^43^, L. Krueger^62^, L. Marouzet^32^, L. Hewitt^32^, L.J. Fleming^4^, M. Kupczyk^8^, M. Ericsson^63^, M. Rahman-Amin^30^, M. Santoninco^26^, M. Sjödin^8^, A. Berton^52^, M. Gerhardsson de Verdier^52^, M. Mikus^64^, M. van de Pol^13^, M. van Geest^52^, M. Gahlemann^65^, Basel, Switzerland^66^, M. Robberechts^16^, M. Szentkereszty^43^, M. Caruso^37^, M.J. Loza^67^, M. Klüglich^35^, M. Kots^68^, M. Rutgers^55^, M. Miralpeix^51^, N. Mores^44^, N. Vissing^47^, N. Rao^69^, N. Fitch^70^, N. Gozzard^71^, N. Lazarinis^39^, N. Adriaens^13^, N. Krug^25^, P.J. Carvalho^4^, P. Söderman^72^, P. Montuschi^44^, P. Chanez^73^, P. Dennison^56^, P. Brinkman^13^, P. Bakke^74^, P. Howarth^75^, P. Nilsson^64^, P. Monk^76^, P. Badorrek^36^, P.-P. Hekking^13^, P. de Boer^55^, P. Powell^77^, R. Sigmund^35^, R. Lutter^13^, R. Hu^3^, R. Middelveld^8^, R. Chaleckis^31^, R. Emma^37^, S. Lone-Latif^13^, S. Meah^4^, S. Valente^44^, S. Walker^30^, S. Pink^32^, S. Masefield^77^, S. Kuo^4^, S. Wagers^70^, S. Naz^8^, S. Williams^48^, S. Hu^4^, S. Hashimoto^13^, S. Reinke^8^, S. Pavlidis^4^, S.J. Fowler^27^, S.J. Wilson^50^, S. Palkonen^79^, S.-E. Dahlén^8^, T. Dekker^13^, T. Geiser^80^, T. Sandström^19^, T. Higgenbottam^81^, U. Nihlen^52^, U. Frey^82^, U. Hoda^83^, V. Hudson^30^, V. Erpenbeck^84^, W. Yu^3^, W. Zetterquist^18^, W. van Aalderen^13^, W. Seibold^35^, X. Yang^4^, X. Hu^3^, Y.-k. Guo^9^, Z. Weiszhart^85^.

^16^MSD, Brussels, Belgium

^17^Royal Brompton and Harefield NHS Foundation Trust, London, UK

^18^Dept of Women’s and Children’s Health and Centre for Allergy Research, Karolinska Institutet, Stockholm, Sweden

^19^Dept of Public Health and Clinical Medicine, Umeå University, Umeå, Sweden

^20^University of Southampton, Southampton, UK

^21^Janssen R&D, High Wycombe, UK

^22^Philips Research Laboratories, Eindhoven, The Netherlands

^23^Lega Italiano Anti Fumo, Catania, Italy

^24^Merck Research Laboratories, Boston, MA, USA

^25^Fraunhofer Institute for Toxicology and Experimental Medicine, Hannover, Germany

^26^University of Rome “Tor Vergata”, Rome, Italy

^27^Centre for Respiratory Medicine and Allergy, Institute of Inflammation and Repair, University of Manchester and University Hospital of South Manchester, Manchester Academic Health Sciences Centre, Manchester, UK

^28^University of Gent, Gent, Belgium

^29^MSD, Kenilworth, NJ, USA

^30^Asthma UK, London, UK

^31^Centre of Allergy Research, Karolinska Institutet, Stockholm, Sweden

^32^NIHR Southampton Respiratory Biomedical Research Unit, Southampton, UK

^33^Respiratory and Allergy Diseases, Genentech, San Francisco, CA, USA

^34^University Hospital Bern, Bern, Switzerland

^35^Boehringer Ingelheim Pharma, Biberach, Germany

^36^Fraunhofer ITEM, Hannover, Germany

^37^Dept of Clinical and Experimental Medicine, University of Catania, Catania, Italy

^38^Respiratory Research Unit, University of Nottingham, Nottingham, UK

^39^Karolinska University Hospital and Karolinska Institutet, Stockholm, Sweden

^40^UK Clinical Operations, GSK, Uxbridge, UK

^41^University Children’s Hospital, Zurich, Switzerland

^42^Janssen R&D, Spring House, PA USA

^43^Semmelweis University, Budapest, Hungary

^44^Università Cattolica del Sacro Cuore, Milan, Italy

^45^NIHR Southampton Respiratory Biomedical Research Unit, Clinical and Experimental Sciences and Human Development and Health, Southampton, UK

^46^II Dept of Internal Medicine, Jagiellonian University Medical College, Krakow, Poland

^47^COPSAC, Copenhagen Prospective Studies on Asthma in Childhood, Herlev and Gentofte Hospital, University of Copenhagen, Copenhagen, Denmark

^48^International Primary Care Respiratory Group, Aberdeen, UK

^49^Karolinska University Hospital, Sweden

^50^Histochemistry Research Unit, Faculty of Medicine, University of Southampton, Southampton, UK

^51^Almirall, Barcelona, Spain

^52^AstraZeneca, Mölndal, Sweden

^53^Royal Brompton and Harefield NHS Foundation Trust, UK

^54^Areteva R&D, Nottingham, UK

^55^Longfonds, Amersfoort, The Netherlands

^56^NIHR Southampton Respiratory Biomedical Research Unit, Clinical and Experimental Sciences, NIHR-Wellcome Trust Clinical Research Facility, Faculty of Medicine, University of Southampton, Southampton, UK

^57^University of Nottingham, Nottingham, UK

^58^CromSource, Stirling UK

^59^Dept of Women’s and Children’s Health, Uppsala University, Sweden

^60^AstraZeneca, Mohlndal, Sweden

^61^Assistance publique des Hôpitaux de Marseille, Clinique des bronches, allergies et sommeil Espace Éthique Méditerranéen, Aix-Marseille Université, Marseille, France

^62^University Children’s Hospital Bern, Bern, Switzerland

^63^Karolinska University Hospital, Stockholm, Sweden

^64^Science for Life Laboratory and The Royal Institute of Technology, Stockholm, Sweden

^65^Boehringer Ingelheim

^66^Janssen R&D, Spring House, PA, USA

^67^Chiesi Pharmaceuticals, Parma, Italy

^68^Janssen R&D, San Diego, CA, USA

^69^BioSci Consulting, Maasmechelen, Belgium

^70^UCB, Slough, UK

^71^Dept of Women’s and Children’s Health, Karolinska Institutet, Stockholm, Sweden

^72^Assistance publique des Hôpitaux de Marseille, Clinique des bronches, allergies et sommeil, Aix-Marseille Université, Marseille, France

^73^Dept of Clinical Science, University of Bergen, Bergen, Norway

^74^NIHR Southampton Respiratory Biomedical Research Unit, Clinical and Experimental Sciences, Southampton, UK

^75^Synairgen Research, Southampton, UK

^76^European Lung Foundation, Sheffield, UK

^77^European Federation of Allergy and Airways Diseases Patient’s Associations, Brussels, Belgium

^78^Dept of Respiratory Medicine, University Hospital Bern, Switzerland

^79^Allergy Therapeutics, Worthing, UK

^80^University Children’s Hospital, Basel, Switzerland

^81^Imperial College, London, UK

^82^Translational Medicine, Respiratory Profiling, Novartis Institutes for Biomedical Research, Basel, Switzerland

^83^Semmelweis University, Budapest, Hungary

^84^University of Southampton, Southampton

## Author Contribution statement

JPRS and PJSkipp wrote the main manuscript text. JPRS prepared all figures. JPRS, FS, PJSkipp & R S-G developed the methodology for Morse clustering in a TDA network. KS and IP processed, integrated and curated gene expression and patient clinical and demographic data. JPRS, JB, MB, IA, KFC, AB, RK, S-ED, CW, JR, CA, BDM, DL, DR, AS, PJSterk, RE, BM, RD, R S-G and PJS planned the investigation and contributed to revising the manuscript.

## Acknowledgments

This paper is presented on behalf of the U-BIOPRED Study Group with input from the U-BIOPRED Patient Input Platform, Ethics Board and Safety Management Board. We thank all the members of each recruiting centre for their dedicated effort, devotion, promptness and care in the recruitment and assessment of the participants in this study. U-BIOPRED is supported through an Innovative Medicines Initiative Joint Undertaking under grant agreement no. 115010, resources of which are composed of financial contribution from the European Union’s Seventh Framework Programme (FP7/2007–2013) and EFPIA companies’ in-kind contribution (www.imi.europa.eu). We would also like to acknowledge help from the IMI funded eTRIKS project (EU Grant Code No.115446).

The U-BIOPRED consortium wishes to acknowledge the help and expertise of the following individuals and groups without whom, the study would not have been possible. Investigators and contributors: Nora Adriaens, Antonios Aliprantis, Kjell Alving, Per Bakke, David Balgoma, Clair Barber, Frédéric Baribaud, Stewart Bates, An Bautmans, Jorge Beleta, Grazyna Bochenek, Joost Brandsma, Armin Braun, Dominic Burg, Leon Carayannopoulos, João Pedro Carvalho da Purificação Rocha, Romanas Chaleckis, Arnaldo D’Amico, Jorge De Alba, Tamara Dekker, Annemiek Dijkhuis, Aleksandra Draper, Rosalia Emma, Magnus Ericsson, Breda Flood, Hector Gallart, Kerry Gove, Neil Gozzard, Lorraine Hewitt, Jens Hohlfeld, Cecile Holweg, Richard Hu, Sile Hu, Juliette Kamphuis, Erika J. Kennington, Dyson Kerry, Hugo Knobel, Johan Kolmert, Maxim Kots, Scott Kuo, Maciej Kupczyk, Bart Lambrecht, Saeeda Lone-Latif, Lisa Marouzet, Jane Martin, Sarah Masefield, Caroline Mathon, Sally Meah, Andrea Meiser, Leanne Metcalf, Montse Miralpeix, Shama Naz, Ben Nicholas, Peter Nilsson, Jörgen Östling, Antonio Pacino, Susanna Palkonen, Stelios Pavlidis, Giorgio Pennazza, Anne Petrén, Sandy Pink, Anthony Postle, Malayka Rahman-Amin, Navin Rao, Lara Ravanetti, Emma Ray, Stacey Reinke, Leanne Reynolds, John Riley, Martine Robberechts, Amanda Roberts, Kirsty Russell, Michael Rutgers, Marco Santoninco, Corinna Schoelch, James P.R. Schofield, Marcus Sjödin, Paul J. Skipp, Barbara Smids, Caroline Smith, Jessica Smith, Doroteya Staykova, Kai Sun, John-Olof Thörngren, Bob Thornton, Jonathan Thorsen, Marianne van de Pol, Marleen van Geest, Anton Vink, Frans Wald, Samantha Walker, Jonathan Ward, Zsoka Weiszhart, Kristiane Wetzel, Craig E. Wheelock, Coen Wiegman, Siân Williams, Susan J. Wilson, Ashley Woodcock, Xian Yang, Elizabeth Yeyasingham.

Partner organisations: Novartis Pharma AG; University of Southampton, Southampton, UK; Academic Medical Centre, University of Amsterdam, Amsterdam, The Netherlands; Imperial College London, London, UK; University of Catania, Catania, Italy; University of Rome ‘Tor Vergata’, Rome, Italy; Hvidore Hospital, Hvidore, Denmark; Jagiellonian Univ. Medi.College, Krakow, Poland; University Hospital, Inselspital, Bern, Switzerland; Semmelweis University, Budapest, Hungary; University of Manchester, Manchester, UK; Université d’Aix-Marseille, Marseille, France; Fraunhofer Institute, Hannover, Germany; University Hospital, Umea, Sweden; Ghent University, Ghent, Belgium; Ctr. Nat. Recherche Scientifique, Villejuif, France; Università Cattolica del Sacro Cuore, Rome, Italy; University Hospital, Copenhagen, Denmark; Karolinska Institutet, Stockholm, Sweden; Nottingham University Hospital, Nottingham, UK; University of Bergen, Bergen, Norway; Netherlands Asthma Foundation, Leusden, NL; European Lung Foundation, Sheffield, UK; Asthma UK, London, UK; European Fed. of Allergy and Airways Diseases Patients’ Associations, Brussels, Belgium; Lega Italiano Anti Fumo, Catania, Italy; International Primary Care Respiratory Group, Aberdeen, Scotland; Philips Research Laboratories, Eindhoven, NL; Synairgen Research Ltd, Southampton, UK; Aerocrine AB, Stockholm, Sweden; BioSci Consulting, Maasmechelen, Belgium; Almirall; AstraZeneca; Boehringer Ingelheim; Chiesi; GlaxoSmithKline; Roche; UCB; Janssen Biologics BV; Amgen NV; Merck Sharp & Dohme Corp.

Third Parties to the project, contributing to the clinical trial: Academic Medical Centre (AMC), Amsterdam (In the U-BIOPRED consortium the legal entity is AMC Medical Research BV (AMR); AMR is a subsidiary of both AMC and the University of Amsterdam; AMC contribute across the U-BIOPRED project); University Hospital Southampton NHS Trust (third party of the University of Southampton and contributor to the U-BIOPRED clinical trial); South Manchester Healthcare Trust (third party to the University of Manchester, South Manchester Healthcare Trust, contributor to the U-BIOPRED clinical trial and to the U-BIOPRED Biobank); Protisvalor Méditerranée SAS (third party to University of the Mediterranean; contributor to the U-BIOPRED clinical trial); Karolinska University Hospital (third party Karolinska Institutet (KI), contributor to the U-BIOPRED clinical trial); Nottingham University Hospital (third party to University of Nottingham, contributor to the U-BIOPRED clinical trial); NIHR-Wellcome Trust Clinical Research Facility.

Members of the ethics board: Jan-Bas Prins, biomedical research, LUMC, the Netherlands; Martina Gahlemann, clinical care, BI, Germany; Luigi Visintin, legal affairs, LIAF, Italy; Hazel Evans, paediatric care, Southampton, UK; Martine Puhl, patient representation (co-chair), NAF, the Netherlands; Lina Buzermaniene, patient representation, EFA, Lithuania; Val Hudson, patient representation, Asthma UK; Laura Bond, patient representation, Asthma UK; Pim de Boer, patient representation and pathobiology, IND; Guy Widdershoven, research ethics, VUMC, the Netherlands; Ralf Sigmund, research methodology and biostatistics, BI, Germany.

The patient input platform: Amanda Roberts, UK; David Supple (chair), UK; Dominique Hamerlijnck, The Netherlands; Jenny Negus, UK; Juliëtte Kamphuis, The Netherlands; Lehanne Sergison, UK; Luigi Visintin, Italy; Pim de Boer (co-chair), The Netherlands; Susanne Onstein, The Netherlands.

Members of the safety monitoring board: William MacNee, clinical care; Renato Bernardini, clinical pharmacology; Louis Bont, paediatric care and infectious diseases; Per-Ake Wecksell, patient representation; Pim de Boer, patient representation and pathobiology (chair); Martina Gahlemann, patient safety advice and clinical care (co-chair); Ralf Sigmund, bio-informatician.

This work was partially funded by the Engineering and Physical Sciences Research Council, UK (EP/N014189: Joining the Dots, from Data to Insight).

